## Supplementary material for "Deep mutational scanning and machine learning reveal structural and molecular rules governing allosteric hotspots in homologous proteins": all supplemental figures

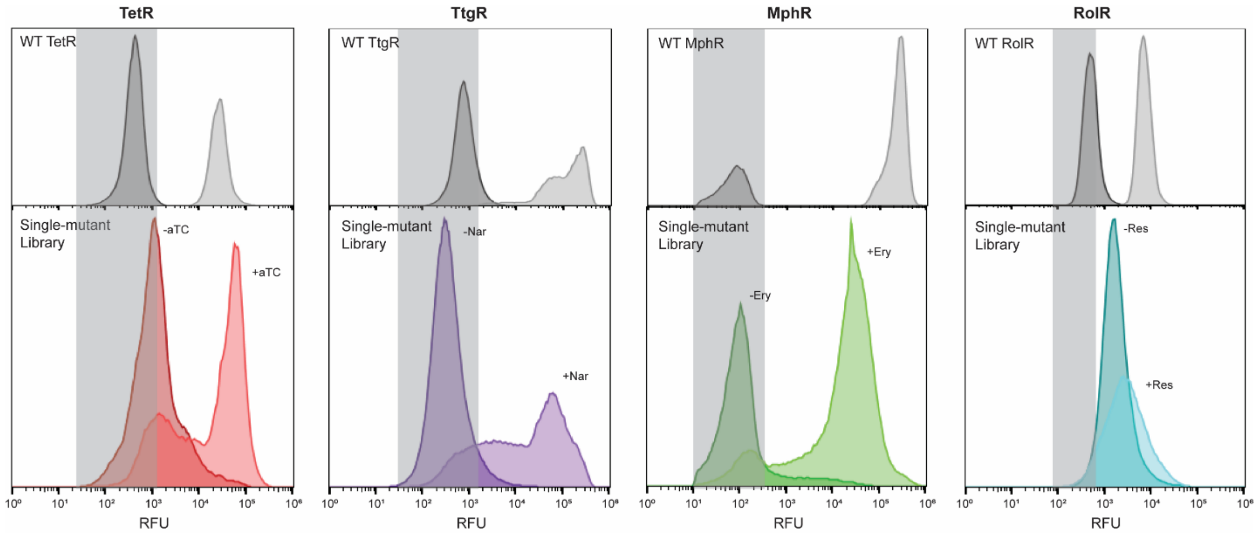


**Figure 1 – figure supplement 1.**

**Sorting scheme identifying inactive variants.** Nonfluorescent cells in the TetR, TtgR, MphR, and RolR single-mutant library were sorted (grey bar) in the presence (light shade) and absence (dark shade) of 1 $\mu$M aTC, 500 $\mu$M Nar, 1 mM Ery, and 7.5 mM Res, respectfully, and sequenced to identify dead variants. Sorting gates were defined by the WT uninduced population for each homolog.

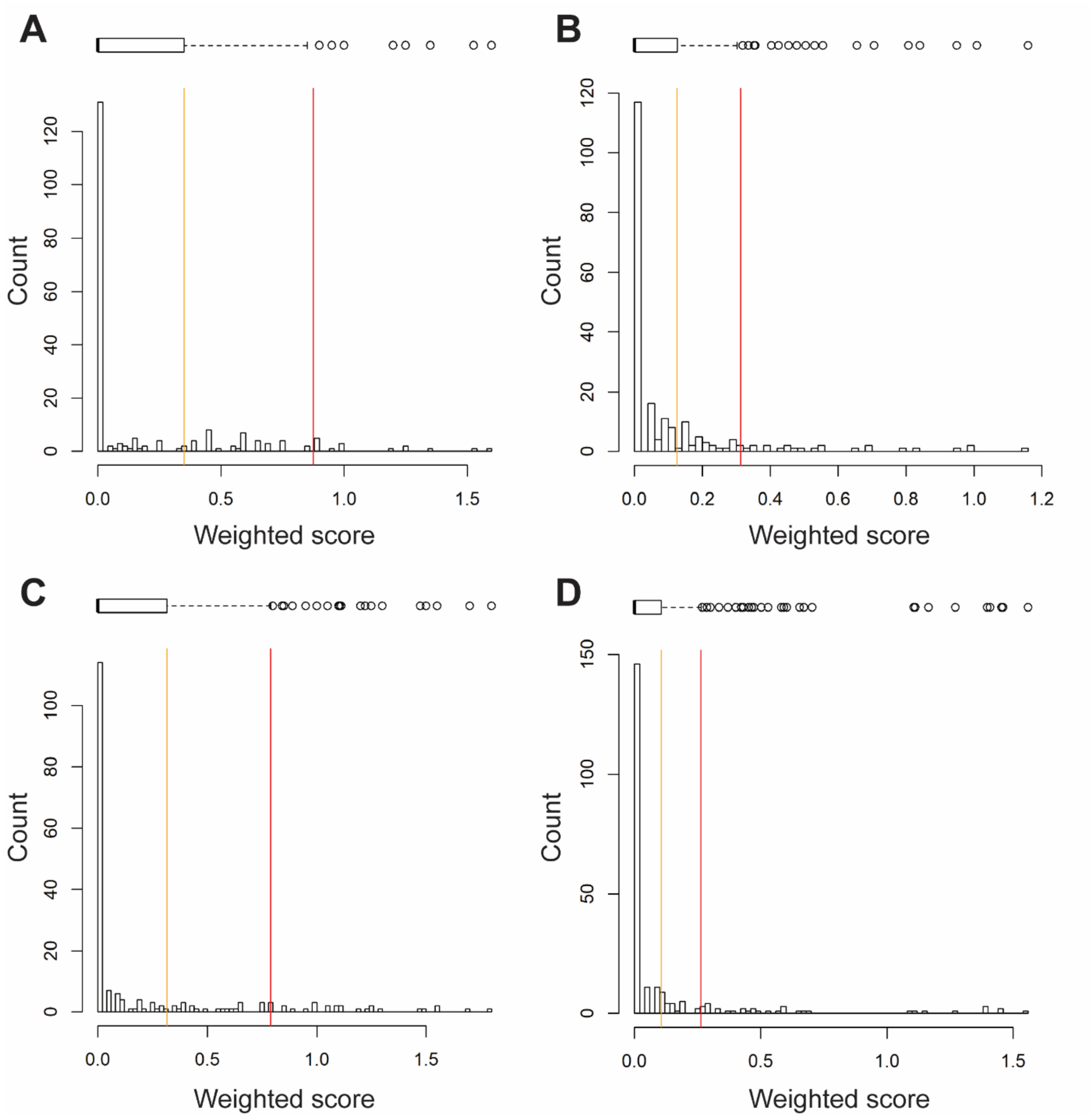


**Figure 1 – figure supplement 2.**

**Histograms of weighted scores and thresholds for identifying hotspots.** The distribution of weighted scores for every position in (A) TetR, (B) TtgR, (C) MphR, and (D) RolR are shown. Box and whisker plots above each histogram illustrate the spread of the data where outliers are shown as circles (red line) and all positions above Q3 (orange line) were designated allosteric hotspots.

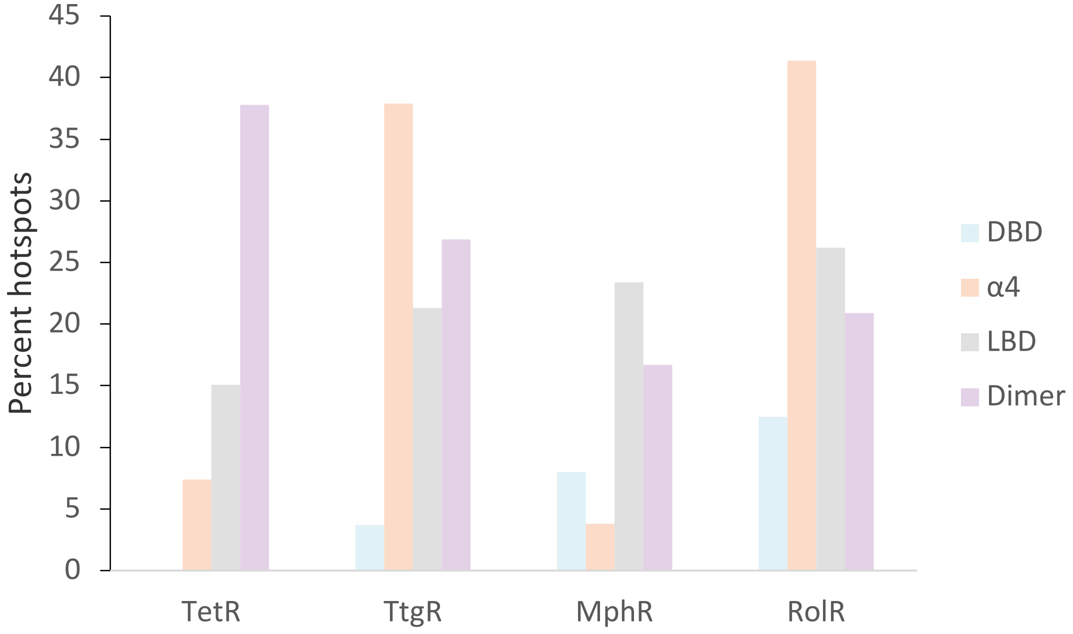


**Figure 1 – figure supplement 3.**
**Distribution of allosteric hotspots in TetR homologs.** The percent of hotspots in the four main structural regions of the TetR homologs. Regions were broken into groups based on the crystal structures of TetR (PDB ID: 4AC0), TtgR (PDB ID: 2UXU), MphR (PDB ID: 3FRQ), and RolR (PDB ID: 3AQT). Potential ligand-binding residues are included in the statistics but are not considered hotspots.

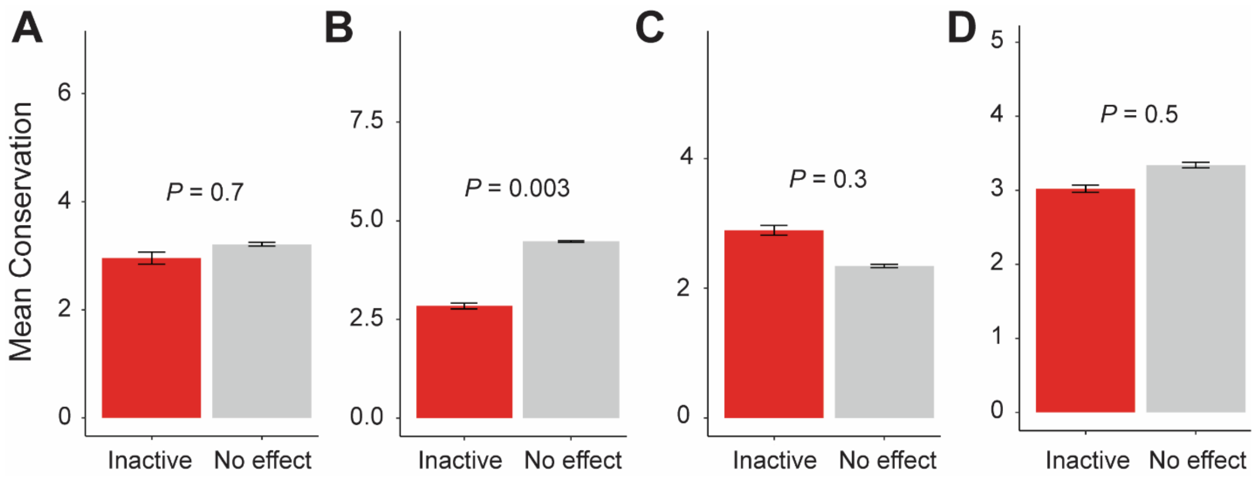


**Figure 1 – figure supplement 4.**

**Conservation of allosteric hotspots**. Average conservation score of all positions considered inactive or having no effect in (A) TetR, (B) TtgR, (C) MphR, and (D) RolR. Data show as mean ± SEM.

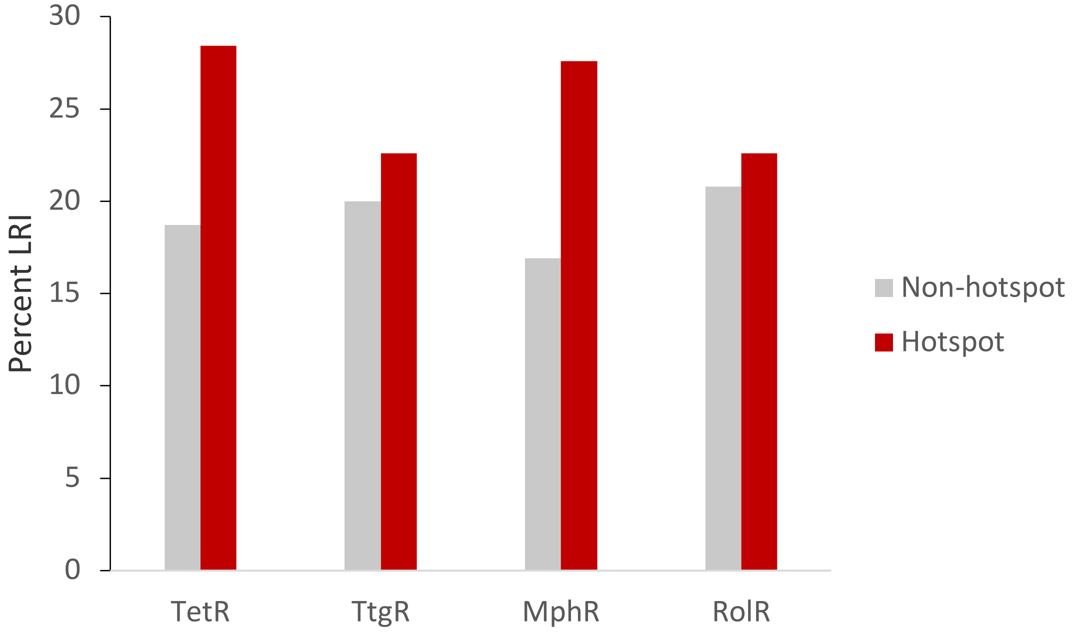


**Figure 2 – figure supplement 1.**
**Hotspot interactions are more likely to be LR than those of non-hotspots.** The percent of hotspot and non-hotspot residues participating in LRIs in each homolog protein.

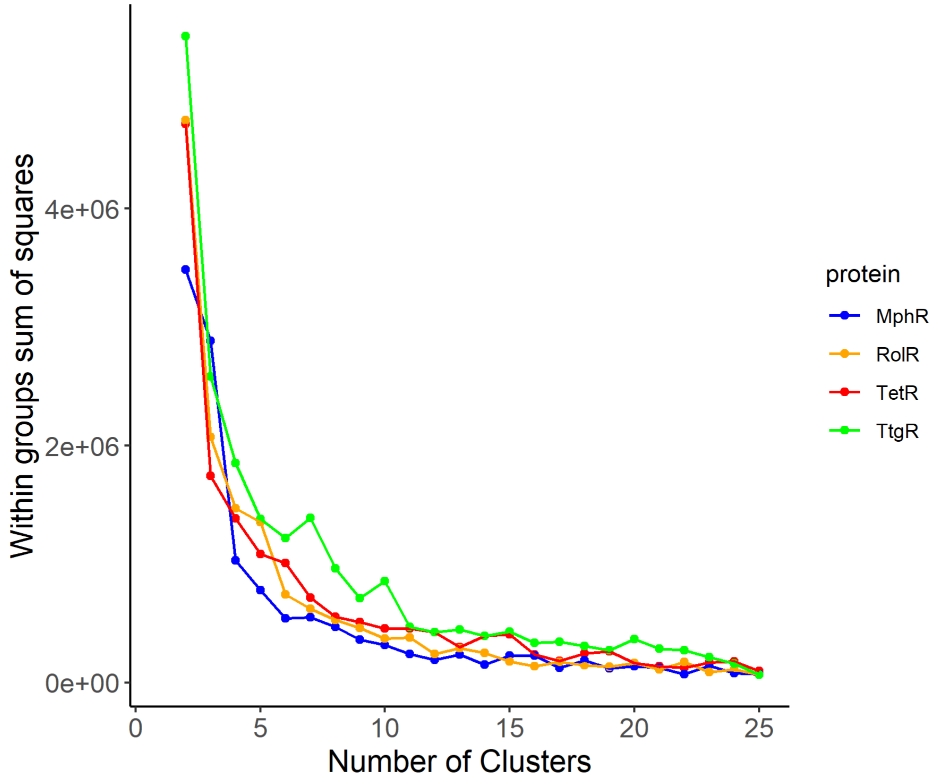


**Figure 2 – figure supplement 2.**
**Elbow method to determine the optimal number of clusters.** The optimal number of clusters to use for the k-means clustering of LRIs in each homolog was determined by iteratively calculating the variance within clusters for 1 to 25 clusters.

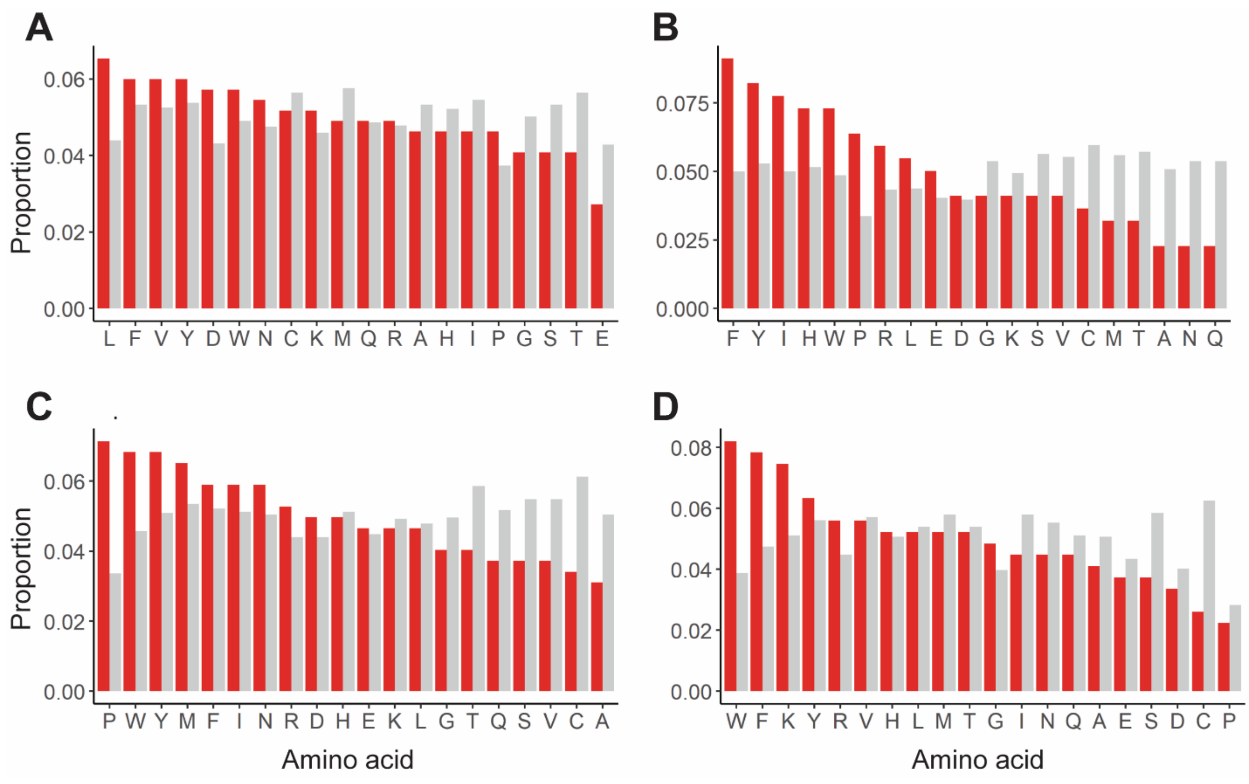


**Figure 3 – figure supplement 1.**
**Enrichment of mutations in allosterically dead or no effect variants.** Mutations in (A) TetR, (B) TtgR, (C) MphR, and (D) RolR were separated based on their effect on protein function, dead (red) or no effect (grey), and the proportion of each of the 20 amino acids within each set calculated to identify enrichments in allosterically dead or neutral variants.

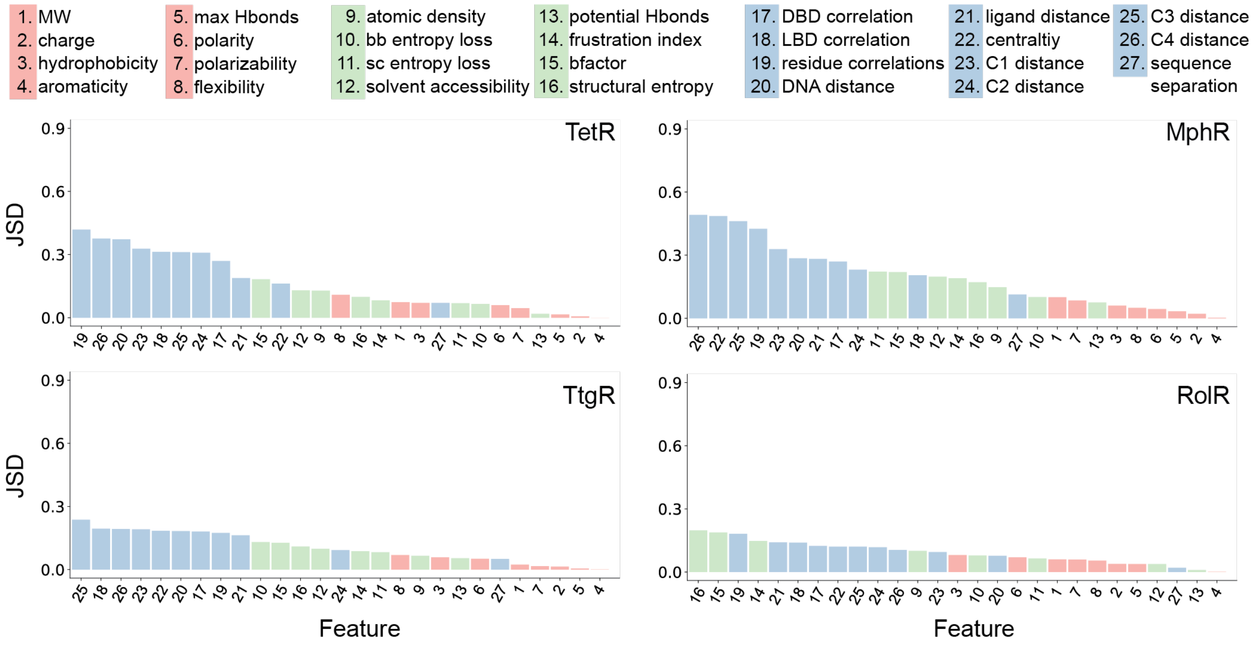


**Figure 4—figure supplement 1.**

**Global features have the highest Jensen-Shannon divergence (JSD).** The full list of 27 features is shown at the top. The JSDs (measure of importance) of the features for each of the four aTFs is shown below. JSD is a measure of similarity between two probability distributions P and Q, which is bound between 0 (P and Q are the same) and 1 (P and Q have no overlap). The larger the JSD, the more different the two distributions are, and thus the features with larger JSDs are more discriminative for hotspot residues. JSD is a symmetrized and smoothed version of the more familiar Kullback-Liebler divergence defined as $JSD(P||Q) = \{ D_{KL}(P||M) + D_{KL}(Q||M) \}/2$, where $M = (P+Q)/2$ is the average of two distributions and $D_{KL}$ is the Kullback-Liebler divergence (KL divergence) which also measures similarity between two distributions. KL divergence is defined as $D_{KL}(P||M) = \sum_x P(x)*\log_2[P(x)/M(x)]$, x are points of the probability space where discrete distributions P and M are defined.

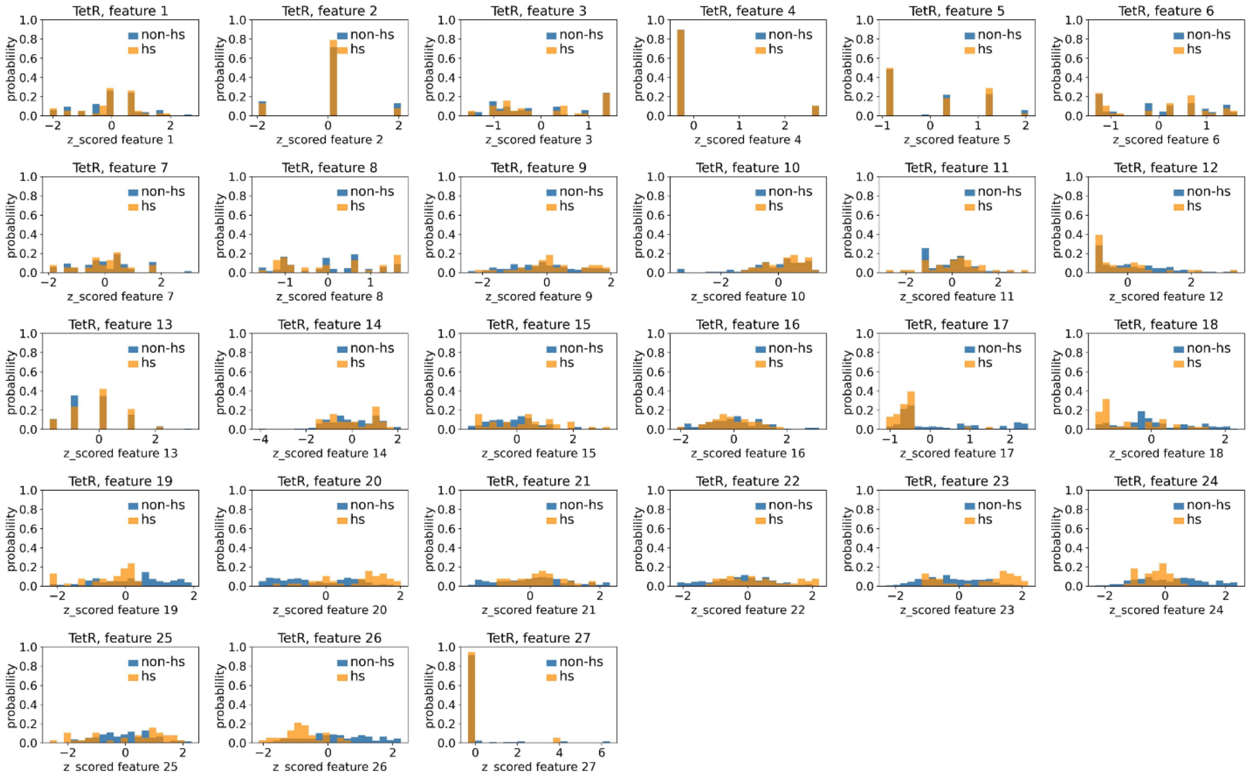

**Figure 4—figure supplement 2.**

**Distributions of TetR's hotspots' and non-hotspots' z-scored feature values for feature 1-27.** The 27 plots correspond to the distributions of TetR's hotspots' (hs) and non-hotspots' (non-hs) z-scored feature values for feature 1-27 as labeled by figure titles. The distributions of hotspots and non-hotspots are normalized by their populations, thus the y axis of the figures are probabilities. Z-scored feature j value of a residue n ($Z_{nj}$), is defined as the difference between its raw feature j value ($R_{nj}$) and the average raw feature j values of all residues (avg_$R_i$), divided by the standard deviation of raw feature j values of all residues, $Z_{nj}$ = ($R_{nj}$ - avg_$R_j$)/std_$R_j$.

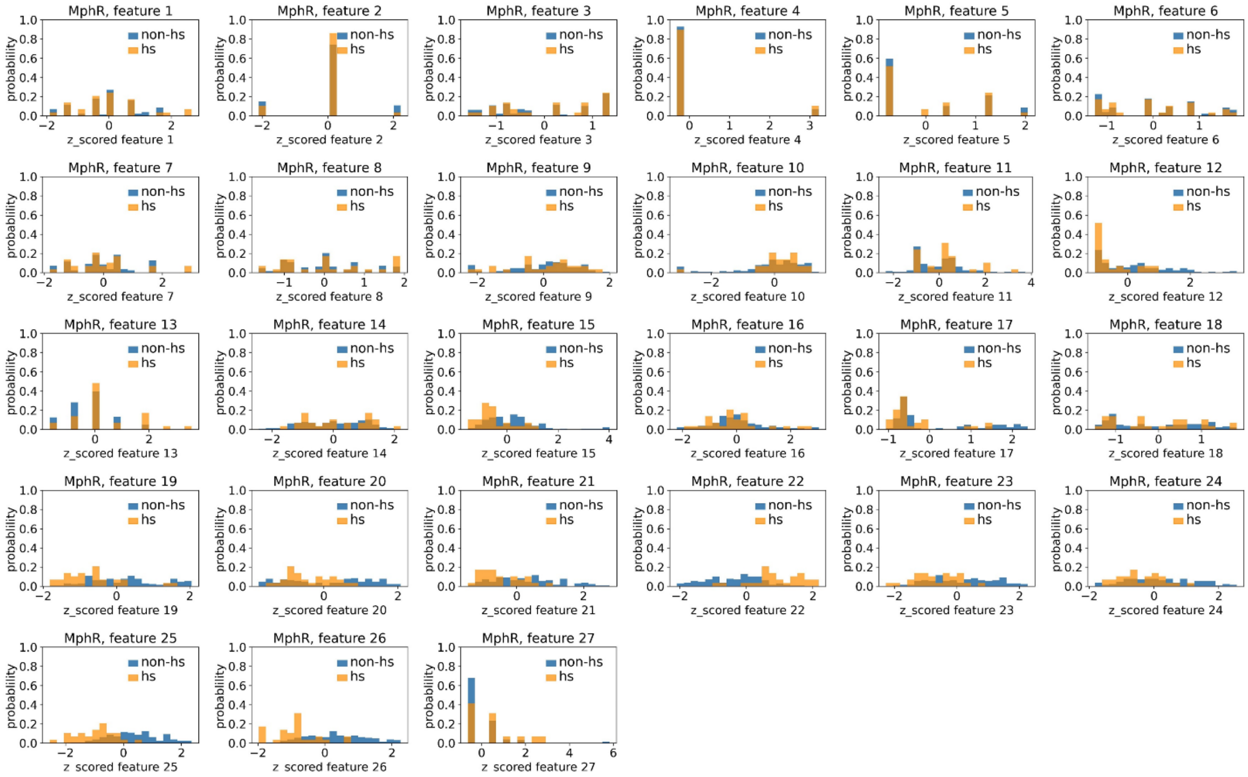


**Figure 4—figure supplement 3.**

**Distributions of MphR's hotspots' and non-hotspots' z-scored feature values for feature 1-27.** The 27 plots correspond to the distributions of MphR's hotspots' (hs) and non-hotspots' (non-hs) z-scored feature values for feature 1-27 as labeled by figure titles. The distributions of hotspots and non-hotspots are normalized by their populations, thus the y axis of the figures are probabilities. Z-scored feature j value of a residue n ($Z_{nj}$), is defined as the difference between its raw feature j value ($R_{nj}$) and the average raw feature j values of all residues ($avg\_R_i$), divided by the standard deviation of raw feature j values of all residues, $Z_{nj} = (R_{nj} - avg\_R_j)/std\_R_j$.

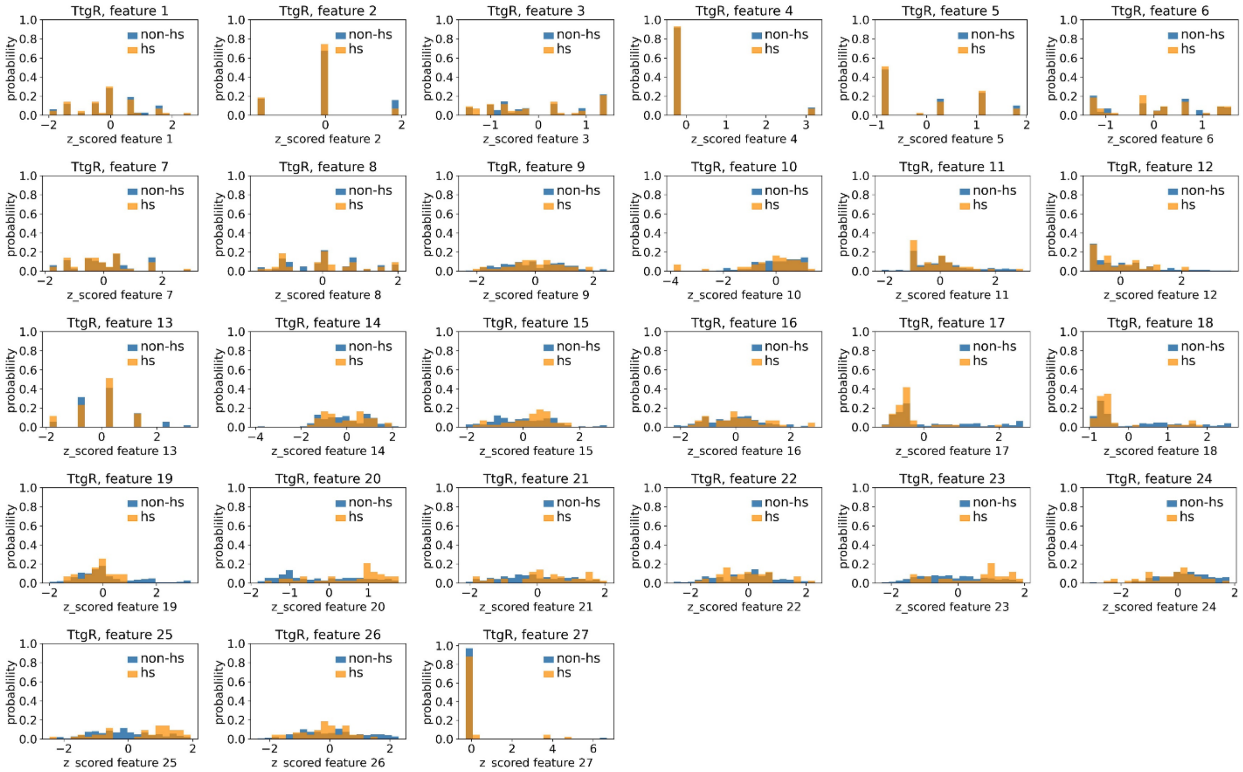


**Figure 4—figure supplement 4.**

**Distributions of TtgR's hotspots' and non-hotspots' z-scored feature values for feature 1-27.** The 27 plots correspond to the distributions of TtgR's hotspots' (hs) and non-hotspots' (non-hs) z-scored feature values for feature 1-27 as labeled by figure titles. The distributions of hotspots and non-hotspots are normalized by their populations, thus the y axis of the figures are probabilities. Z-scored feature j value of a residue n ($Z_{nj}$), is defined as the difference between its raw feature j value ($R_{nj}$) and the average raw feature j values of all residues (avg_$R_i$), divided by the standard deviation of raw feature j values of all residues, $Z_{nj}$ = ($R_{nj}$ - avg_$R_j$)/std_$R_j$.

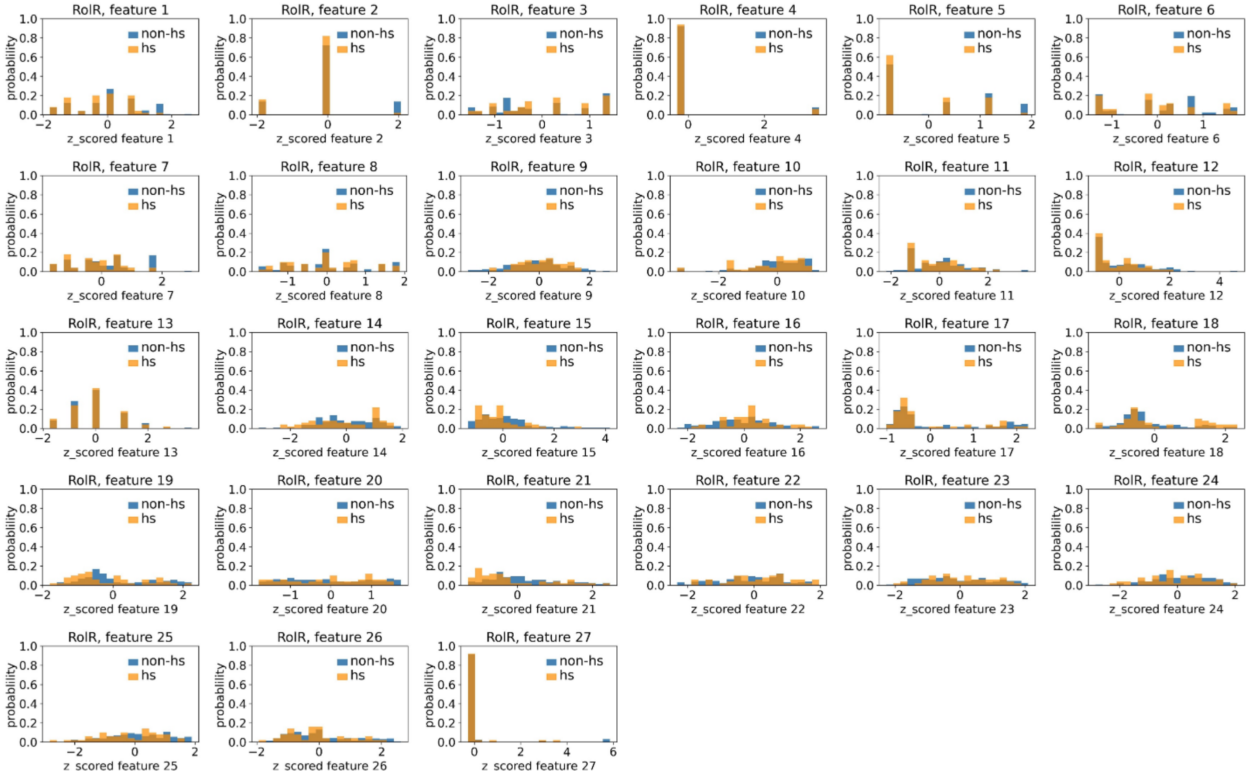


**Figure 4—figure supplement 5.**

**Distributions of RoIR's hotspots' and non-hotspots' z-scored feature values for feature 1-27.** The 27 plots correspond to the distributions of RoIR's hotspots' (hs) and non-hotspots' (non-hs) z-scored feature values for feature 1-27 as labeled by figure titles. The distributions of hotspots and non-hotspots are normalized by their populations, thus the y axis of the figures are probabilities. Z-scored feature j value of a residue n ($Z_{nj}$), is defined as the difference between its raw feature j value ($R_{nj}$) and the average raw feature j values of all residues ($avg\_R_i$), divided by the standard deviation of raw feature j values of all residues, $Z_{nj} = (R_{nj} - avg\_R_j)/std\_R_j$.

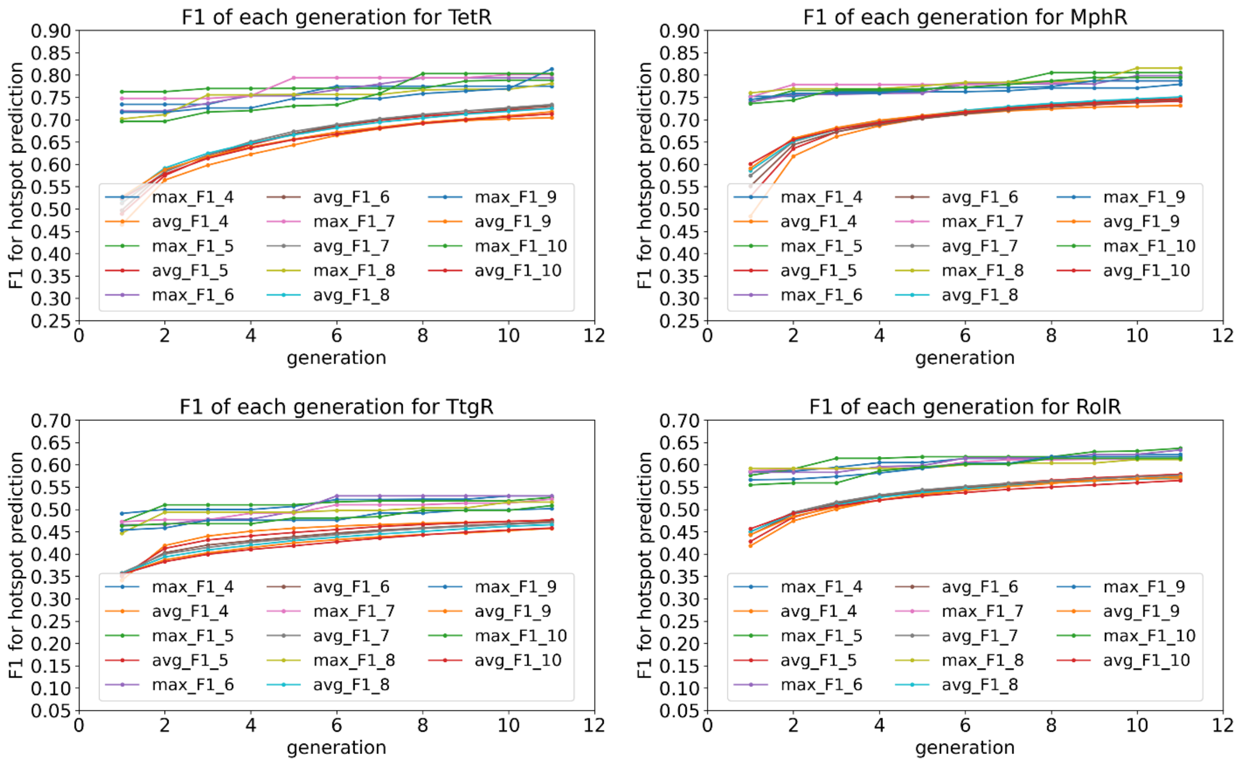


**Figure 4—figure supplement 6.**

**Average and best F1 scores of 4-10 feature combinations converge after 10 generations in the genetic algorithm feature selection.** The plots show the average and best F1 scores for 4-10 feature combinations as a function of generation in the genetic algorithm feature selection for the four homologous aTFs as labeled by figure titles.

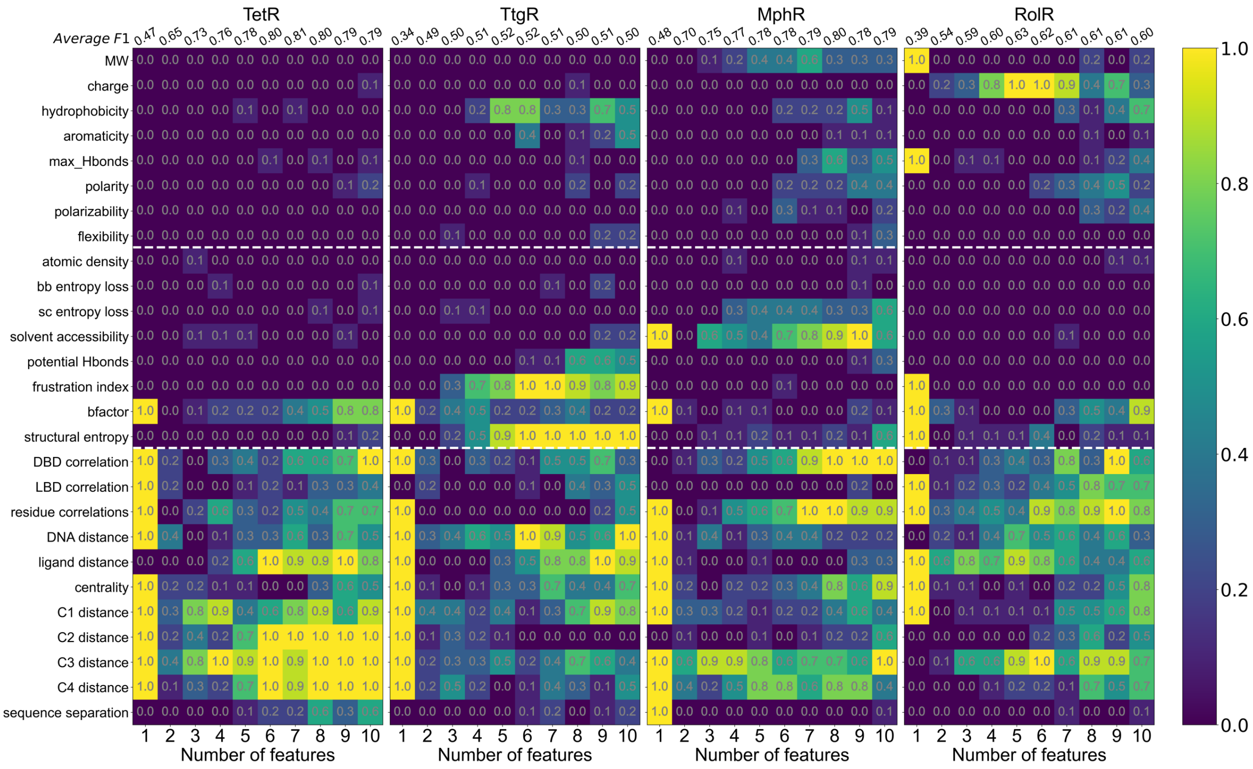


**Figure 4—figure supplement 7.**

**Machine learning identifies structural and molecular features that differentiate allosteric hotspots.** Frequency of appearance of the 27 features in the top ten 1- to 10-feature-combinations ranked by F1 score for each protein (labeled on top). Row 2-28 corresponds to feature 1-27, row 1 is the average F1 score of the top 10 1- to 10-feature combinations. This is the same data as that of Figure 4B with all the frequencies specified in the heat map.

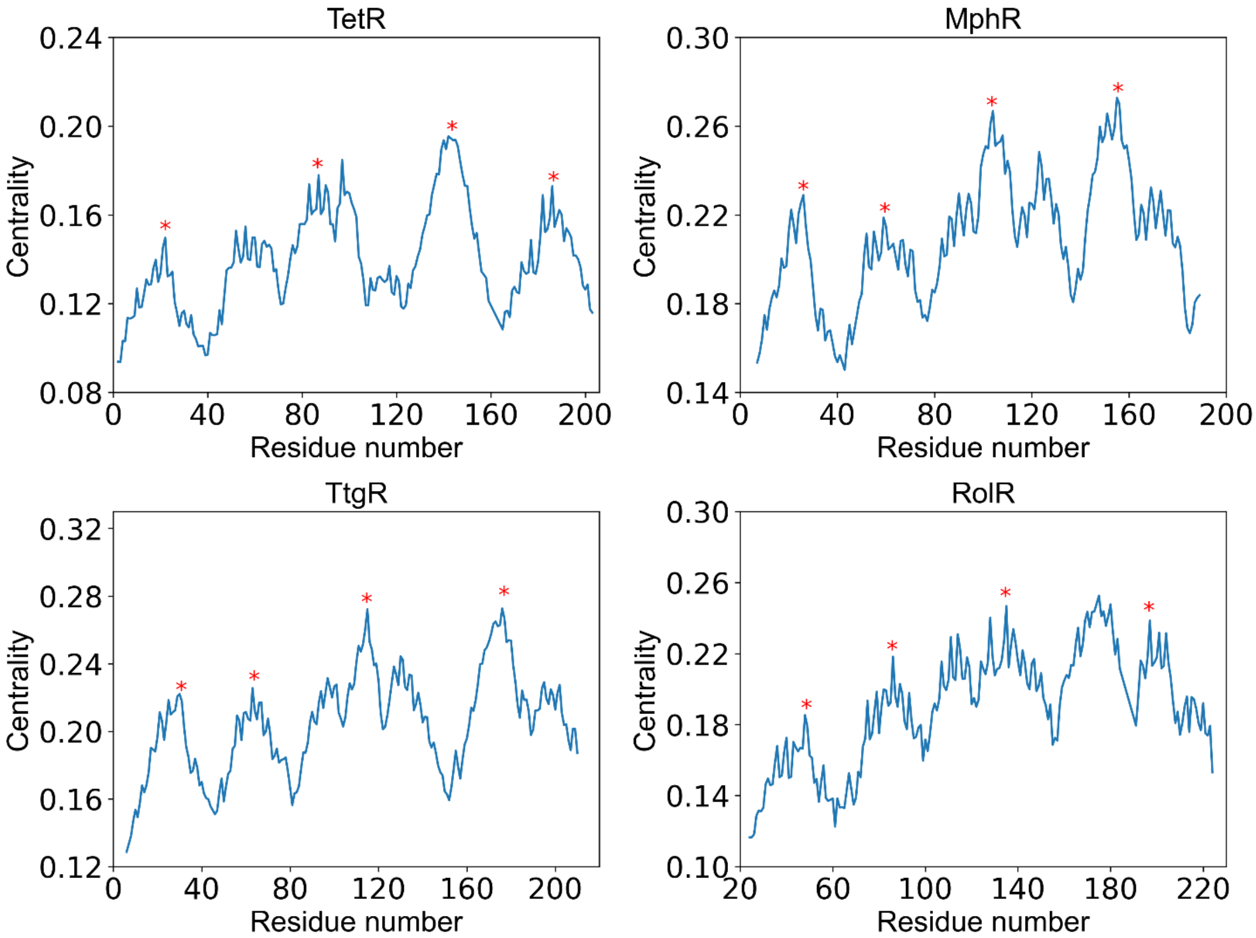


**Figure 4—figure supplement 8.**

**Positions of Centrality peaks.** Plots of centrality against residue number of each protein (labeled by the title), with the four red stars label the positions of centrality peaks 1-4 from left to right. The centrality peaks are identified as positions of highest centrality within local sequence while maintaining distances between centrality peaks as large as possible. The centrality peaks 1-4 are located at residue 22, 83, 150, 193 for TetR; residue 26, 59, 104, 155 for MphR; residue 30, 71, 122, 186 for TtgR; and residue 48, 84, 133, 191 for RolR.

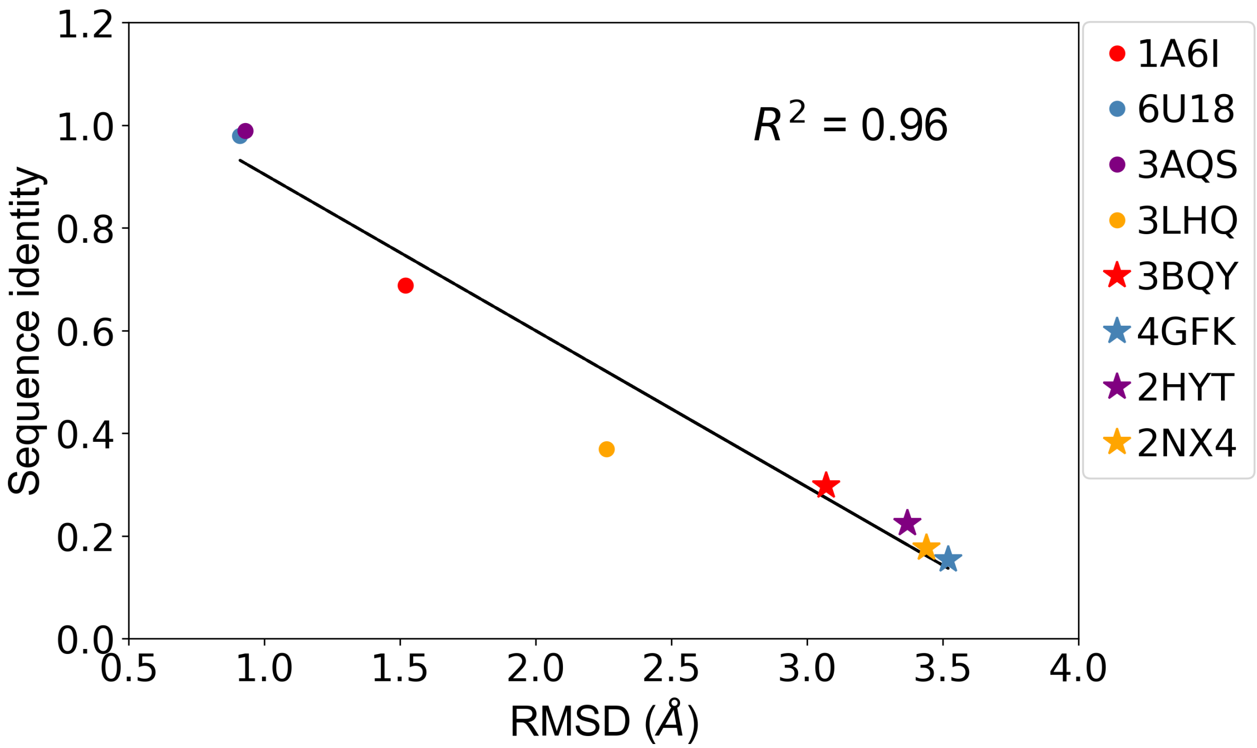


**Figure 6—figure supplement 1.**

**Sequence identity and RMSD between template protein and target protein are anti-correlated.** Correlation of sequence identity and RMSD between the four aTFs and their corresponding templates used in generating homology models. R squared shows the coefficient of determination of the corresponding linear regression. (Red: templates for TetR; Blue: templates for MphR; Purple: templates for RolR; Orange: templates for TtgR.)
